# Broaden the application of *Yarrowia Lipolytica* synthetic biology tools to explore the potential of Yarrowia clade biodiversity

**DOI:** 10.1101/2023.06.05.543681

**Authors:** Young-Kyoung Park, Tristan Rossignol

**Affiliations:** Université Paris-Saclay, INRAE, AgroParisTech, Micalis Institute, 78350, Jouy-en-Josas, France

## Abstract

Yeasts have established themselves as prominent microbial cell factories, and the availability of synthetic biology tools has led to breakthroughs in the rapid development of industrial chassis strains. The selection of a suitable microbial host is critical in metabolic engineering applications, but it has been largely limited to a few well-defined strains. However, there is growing consideration for evaluating strain diversity, as a wide range of specific traits and phenotypes have been reported even within a specific yeast genus or species. Moreover, with the advent of synthetic biology tools, non-type strains can now be easily and swiftly reshaped.

The yeast *Yarrowia lipolytica* has been extensively studied for various applications such as fuels, chemicals, and food. Additionally, other members of the Yarrowia clade are currently being evaluated for their industrial potential. In this study, we demonstrate the versatility of synthetic biology tools originally developed for *Y. lipolytica* by repurposing them for engineering other yeasts belonging to the Yarrowia clade. Leveraging the GoldenGate *Y. lipolytica* tool kit, we successfully expressed fluorescent proteins as well as the carotenoid pathway in at least six members of the clade, serving as proof of concept.

This research lays the foundation for conducting more comprehensive investigations into the under-characterized strains within the Yarrowia clade and exploring their potential applications in biotechnology.

## Introduction

Bio-based products through the microbial fermentation process to replace petroleum-based chemical production are extensively studied as an essential way to develop sustainable and environmentally friendly bioproduction. A key point for microbial cell factory success is the choice of the chassis microorganism for its metabolite production and its metabolic rewiring capacity as well as the availability of synthetic biology tools. Yeasts are particularly attractive eukaryotic production hosts with their capacity to grow on a broad variety of renewable carbon sources, to produce complex target molecules, and their well-established scale-up process.

Apart from *Saccharomyces cerevisiae*, non-conventional yeasts have been investigated due to their particular industrial traits. Among them, *Yarrowia lipolytica* has been extensively used and studied for many applications from fuels and chemicals to foods. This yeast is genetically amenable, can grow on a large range of substrates, has the GRAS status, and can produce high yields of various biomolecules, in particular lipids and organic acids, (Park and Ledesma-Amaro, 2023). Target molecules produced by this yeast have been recently expanded to plant-based derivative products like terpenes (Zhang, et al., 2022). This continuously growing interest in using this yeast as an industrial chassishas led to an exponential advancements in the development of synthetic biology tools including fast and modular cloning systems and CRISPR/Cas9 methods (Larroude, et al., 2018).

Yi et *al*. recently reviewed the importance of considering strain diversity for establishing microbial cell factories by highlighting the fact that a wide range of strain variations already exists even within a specific yeast genus and species (Yi and Alper, 2022). An illustrative example has been provided by a primary screening of the potential producer of triacetic acid lactone by expressing a 2-pyrone synthase in 13 industrial *S. cerevisiae* strains with different genetic background (Saunders, et al., 2015). The production levels varied up to 63-fold among different strains, highlighting the importance of considering the strain variation for optimal microbial cell factories. Moreover, basal production is not always indicative of the rewiring potential (Yi and Alper, 2022), and phenotypes and physiology differ for non-native metabolites. Therefore, heterologous expression must be screened in various backgrounds to evaluate the best strains/species for further genetic engineering. This emphasizes the growing importance of considering strain-level characteristics when selecting host microorganism. However, this requires extensive study of variants or extensive screening with the possible requirements of genetic tools adaptations to each strain.

Most studies focus on a limited set of industrial or type strains and rarely extend to other strains or related species, limiting the knowledge on the physiologic and metabolic state of such species or strains. Nevertheless, for some traits of interest like lipid production, physiological data of clade or set of strains have been explored. The diversity of closely related species in the Yarrowia clade has been recently investigated through comparative studies of their oleaginous properties (Kurtzman, 2011). These species, *Y. lipolytica, Yarrowia alimentaria, Yarrowia deformans, Yarrowia galli, Candida hispaniensis, Yarrowia hollandica, Yarrowia oslonensis, Yarrowia phangngensis* and *Yarrowia yakushimensis* exhibited diverse salt stress tolerance, optimal temperature for growth (Groenewald, et al., 2014) and maximum lipid content, with significant differences ranging from 30% of cell dry weight in *C. oslonensis* to 67% in *C. hispaniensis* (Michely, et al., 2013). In the past decade, the clade has grown up to 15 potential species including now *Yarrowia keelungensis, Yarrowia divulgata, Yarrowia porcina, Yarrowia bubula, Yarrowia brassicae* and *Yarrowia parophonii*, (Quarterman, et al., 2017, Chang, et al., 2013, Péter, et al., 2019). Quarterman et *al*. investigated the biomass and lipid production of 13 species of the Yarrowia clade by using a nondetoxified acid-pretreated switchgrass hydrolysate as a feedstock (Quarterman, et al., 2017) with further characterization of a subset of them for inhibitor tolerance. This study highlighted the diversity in terms of behaviour, production and resistance, with for example, *Y. hollandica* and *Y. phangngensis* reached higher cell biomass and lipid titer compared to *Y. lipolytica* W29 in these conditions. In other studies, up to thirteen of these yeast species from the Yarrowia clade were screened for erythritol, arabitol mannitol, citric acid, lipids and protein production (Rakicka, et al., 2016, Rakicka-Pustułka, et al., 2021). In particular, *Y. divulgata* and *Y. oslonensis* were identified as particularly robust producers of polyol compared to *Y. lipolytica* when grown on glycerol (Rakicka-Pustułka, et al., 2021). Protease production has been evaluated within the Yarrowia clade has confirmed the potential of certain Yarrowia species to secrete significantly high amounts of proteases and revealed the influence of culture media when benchmarking species for a particular trait (Ciurko, et al., 2023).

Therefore, the physiological aspect and diversity of the Yarrowia clade are now well-documented, highlighting the variation in technological traits among different species, which can outperform *Y. lipolytica*. Following this, genetic engineering of the other species of the clade starts to be considered. A first transformation method for the most distant species *C. hispaniensis* allows using sucrose as a carbon source by expressing the *S. cerevisiae* invertase as a marker and a heterologous enzyme. *C. hispaniensis* is resistant to antibiotics like gentamycin, nourseothricin, and hygromycin, which are typically employed as selective markers in yeast genetic engineering. This limits the development of genetic tools in this species (Morin, et al., 2020). Moreover, this yeast is refractory to the standard transformation method and requires the use of biolistic approach for transformation (Morin, et al., 2020). Similarly, *Y. phangngensis* has been transformed for the first time by using genetic tools and the standard transformation method developed for *Y. lipolytica* with hygromycin and zeocin as markers for selections (Quarterman, et al., 2018). The recycling of *Y. lipolytica* tools in this species is shows possibility for more complex genetic engineering. However, it is worth noting that the hygromycin concentration required for selection in *Y. phangngensis* needs to be 3 to 5 times higher compared to *Y. lipolytica* transformants selection. It opens the way for reusing the existed tools or developing the standardized tools in non-conventional and non-type organisms, allowing rapid screening of numerous potential hosts for diverse applications.

Here we evaluated the functionality of a GoldenGate modular synthetic biology tool developed for *Y. lipolytica* (Larroude, et al., 2019) in 6 other species from the Yarrowia clade by using two different selection markers and the expression of a fluorescent protein as a validation system with a fast transformation method. As a proof of concept, the genes in the carotenoid pathway were expressed in 5 of the 6 species.

## Results

### Evaluation of selection markers

The GoldenGate tool kit developed for *Y. lipolytica* contains 2 dominant selection markers, hygromycin and nourseothricin, for selecting transformants. This toolkit enables the expression of up to three transcription units in a single transformation (Larroude, et al., 2019). The minimum concentration of hygromycin and nourseothricin required for growth inhibition was first evaluated on 8 strains of the Yarrowia clade (*Y. lipolytica, Y. alimentaria, Y. deformans, Y. galli, Y. hollandica, Y. oslonensis, Y. phangngensis* and *Y. yakushimensis*) identified in Michely et *al*. (Michely, et al., 2013). All the strains used in this study are listed in Table 1. *C. hispaniensis* was excluded due to its previously established resistance to the major antibiotics (Morin, et al., 2020) and the antibiotics available in the *Y. lipolytica* GoldenGate toolkit would not be effective for this species. The growth of all the strains were inhibited with 200μg/ml hygromycin or 500μg/ml nourseothricin except for *Y. phangngensis* which required higher concentration already demonstrated (Quarterman, et al., 2018). In our hands, we had residual growth of *Y. phangngensis* with 500 μg/mL of hygromycin and 1mg/mL of nourseothricin. Therefore, this specie was not selected thereafter for evaluation of transformability with the *Y. lipolytica* GoldenGate vectors.

**Table1.**
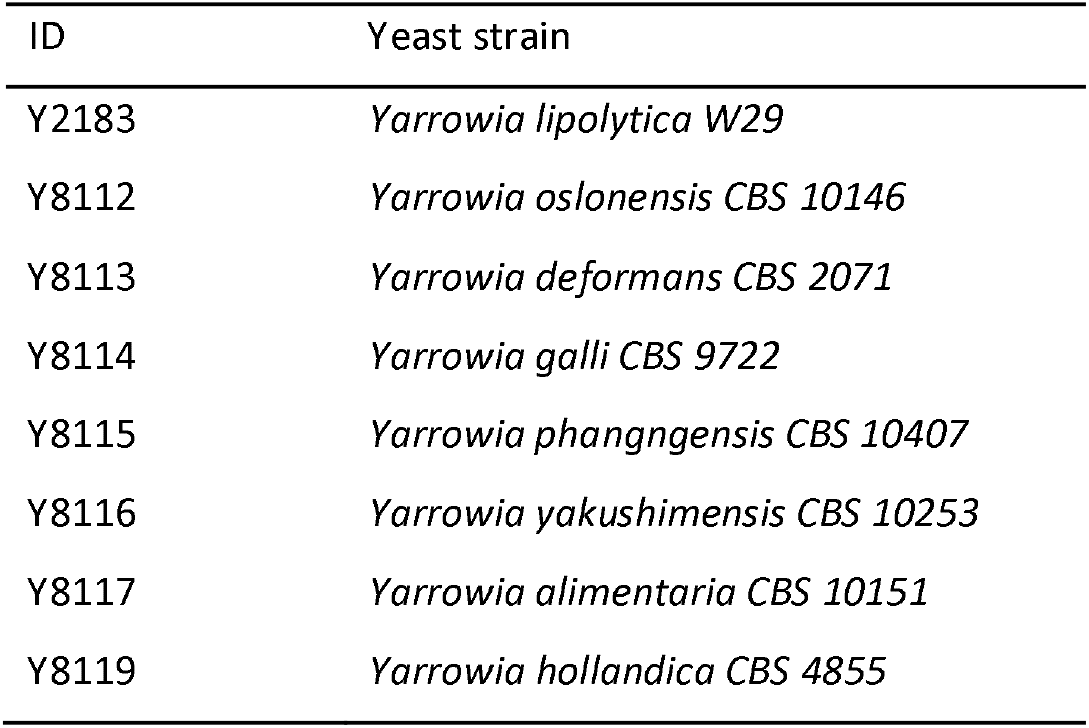
List of wild-type yeast strains of the Yarrowia clade used in this study.

| ID | Yeast strain |
| --- | --- |
| Y2183 | *Yarrowia lipolytica* W29 |
| Y8112 | *Yarrowia oslonensis* CBS 10146 |
| Y8113 | *Yarrowia deformans* CBS 2071 |
| Y8114 | *Yarrowia galli* CBS 9722 |
| Y8115 | *Yarrowia phangngensis* CBS 10407 |
| Y8116 | *Yarrowia yakushimensis* CBS 10253 |
| Y8117 | *Yarrowia alimentaria* CBS 10151 |
| Y8119 | *Yarrowia hollandica* CBS 4855 |

### Evaluation of transformation capacity

To evaluate the transformation of each strain, we used integrative expression vectors assembled using the GoldenGate method containing the expression cassette of the Redstar2 fluorescent protein with either the selective marker hygromycin or nourseothricin, and ZETA sequences that are known for their random integration in the *Y. lipolytica* genome (see material and methods). All the building block have been optimized for *Y. lipolytica* and the details are described in Larroude et *al*. (Larroude, et al., 2019). The two genes present in each construct, the gene coding for the Redstar2 and the gene coding for antibiotic resistance, are under the *Y. lipolytica* pTEF promoter. Therefore, we first evaluated the level of homology between the pTEF promoter sequences of the different strains of the clade. The *Y. lipolytica* pTEF promoter was used to retrieve promoter ortholog from Whole Genome Sequences of the other species. All sequences retrieved are available in supplementary file 1. Sequence alignment showed a strong homology, particularly in the 220 base pair region proximal to the start codon, which suggest the potential for cross-species functionality of the promoter. *Y alimantaria* appears the most distant and is missing a homology region in the distal part of the promoter (supplementary file 2).

The transformation of seven species including *Y. lipolytica* as a positive control were carried out by using a simple and rapid method with a yeast transformation kit (see material and methods section), in line with the objectives of rapid screening of different host without specific transformation setup. Both plasmids expressing the Redstar2 with hygromycin or nourseothricin were tested. We were able to obtain from 1 to around 50 clones for all strains in one transformation experiment except for *Y. alimentaria* for which no clones were obtained even after several attempts and the extended incubation for more than a week which usually helps potential transformants to recover. The relatively lower promoter sequence homology between *C. alimentaria* and *Y. lipolytica*, compared to other species within the clade, might explain the poor expression of selective marker gene resulting in unsuccessful transformation. Individual clones from all transformants were then evaluated for their fluorescence in a microtiter plate reader. All the strains constructed are listed in supplementary Table 1.

All the clones tested were expressing the Redstar2 fluorescent protein with some variation in intensity depending on the species and marker. As the genomic integration takes place randomly, we cannot formally compare the level of fluorescence as genome integration locus may influence the level of heterologous expression. *Y. lipolytica* and *Y. galli* are the species expressing the highest level of fluorescence in average with both markers while *Y. oslonensis* is the being the lowest (Fig. 1).

**Fig. 1.**
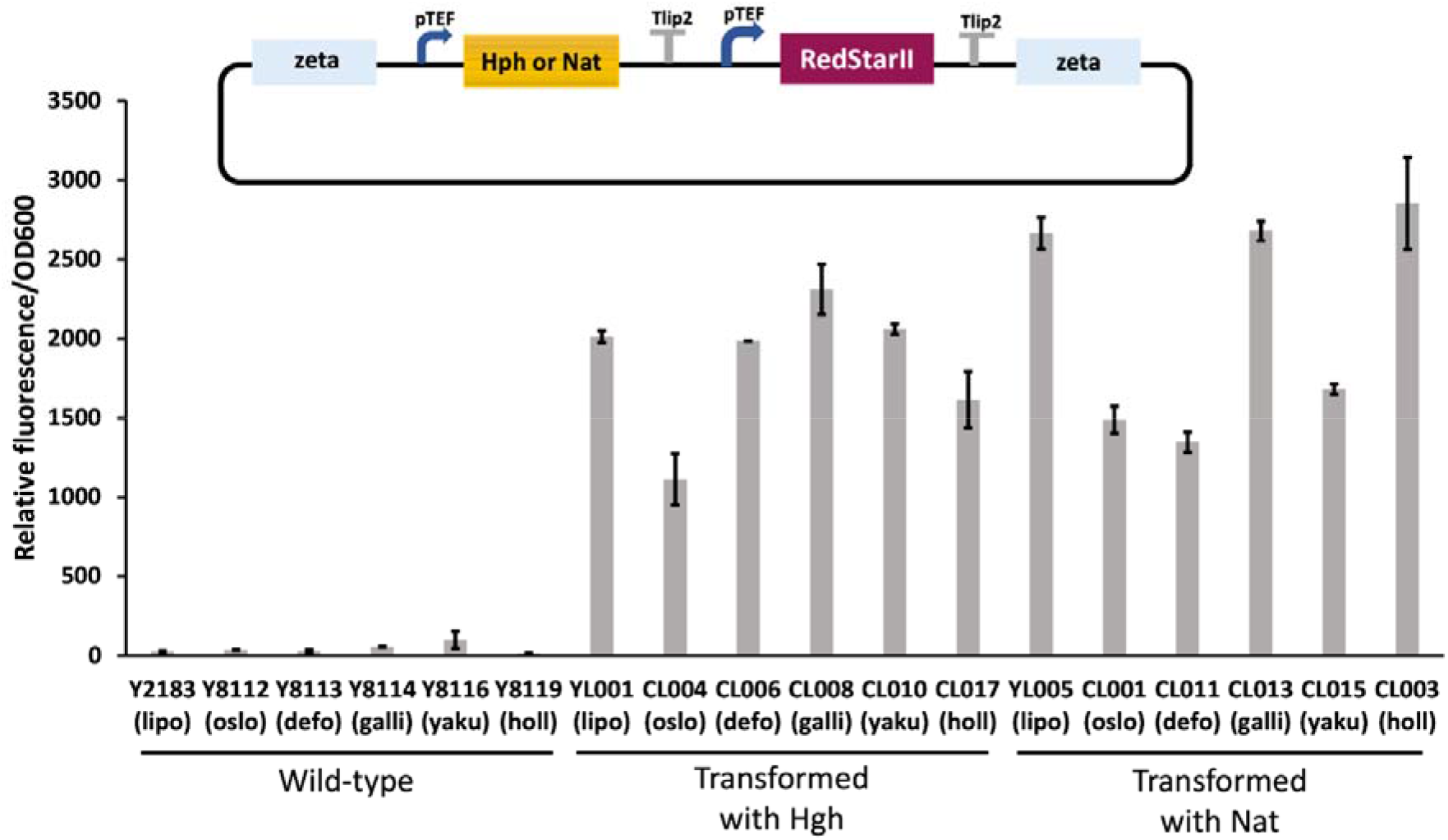
Relative fluorescence reported to OD for wild-type strains and each transformant expressing the Redstar2 protein after 72h into 96-well microplates. Cultures were performed at least in duplicate. The upper illustration correspond to schematic draw of the vector used for transformation. Hgh (vector Hygromycin –pTEF-Redstar2); Nat (vector Nourseothricin –pTEF-Redstar2). lipo (*Y. lipolytica*); oslo (*Y. oslonensis*); defo (*Y. deformans*); galli (*Y. galli*); yaku (*Y. yakushimensis*); holl (*Y. hollandica*). All the strains are listed in supplementary Table 1.

Those differences are not due to the variation in fitness between species as they present a similar growth rate (Fig. 2).

**Fig. 2.**
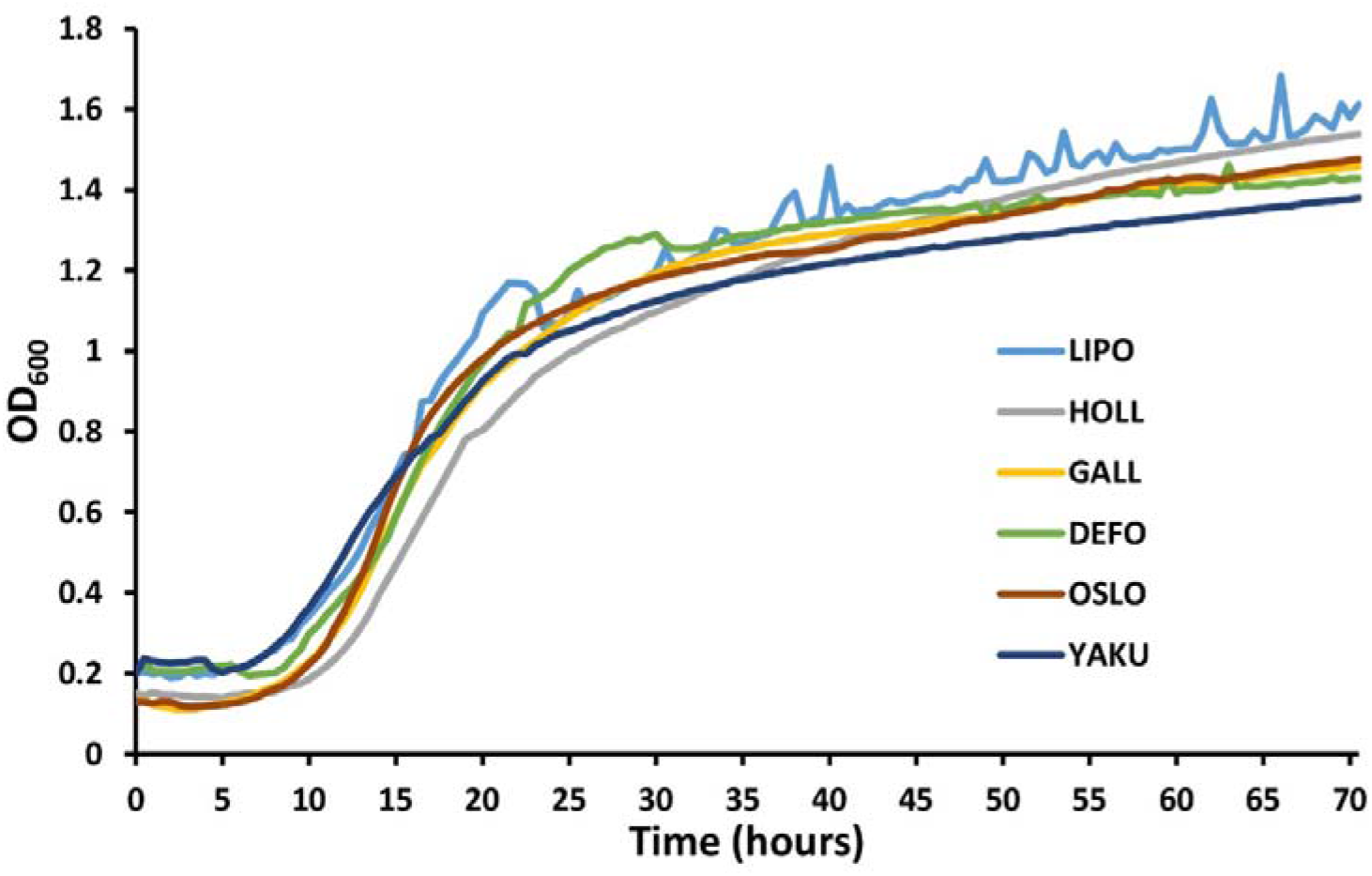
Representative growth curves in 96 well plates for each species. The growth analysis was performed using a microtiter plate reader with a reading every 30 min for 72 h. LIPO (*Y. lipolytica*); OSLO (*Y. oslonensis*); DEFO (*Y. deformans*); GALL (*Y. galli*); YAKU (*Y. yakushimensis*); HOLL (*Y. hollandica*).

### Heterologous pathway expression

The evaluation was pushed forward by the attempt to express a heterologous pathway already established in *Y. lipolytica* allowing the production of carotenoids (Larroude, et al., 2022) with a plasmid containing three transcription units (see material and methods section). This pathway is regularly used as a proof of concept as it allows rapid screening of positive clones with the orange colour. This heterologous pathway-expressing cassette was transformed into seven species, respectively.. As we have seen from the transformation of Redstar2 gene expression cassette, we were not able to obtain transformant *for Y. alimentaria*. We obtained the orange transformants for all the others except for *Y. yakushimensis* (Fig. 3). All the transformants are listed in supplementary Table 1. Again, we got slight differences in colour intensity with *Y. hollandica* having the lowest intensity (Fig. 3). The level of carotenoid could either be due to differences in heterologous expression or metabolic capacity between species for this particular pathway or locus used for the integration even if multiple clones had the same trends in colour intensity. The fact that we have different trends between fluorescent protein expression and carotenoid pathway expression for the same strain is in favour of a metabolic capacity variability.

**Fig. 3.**
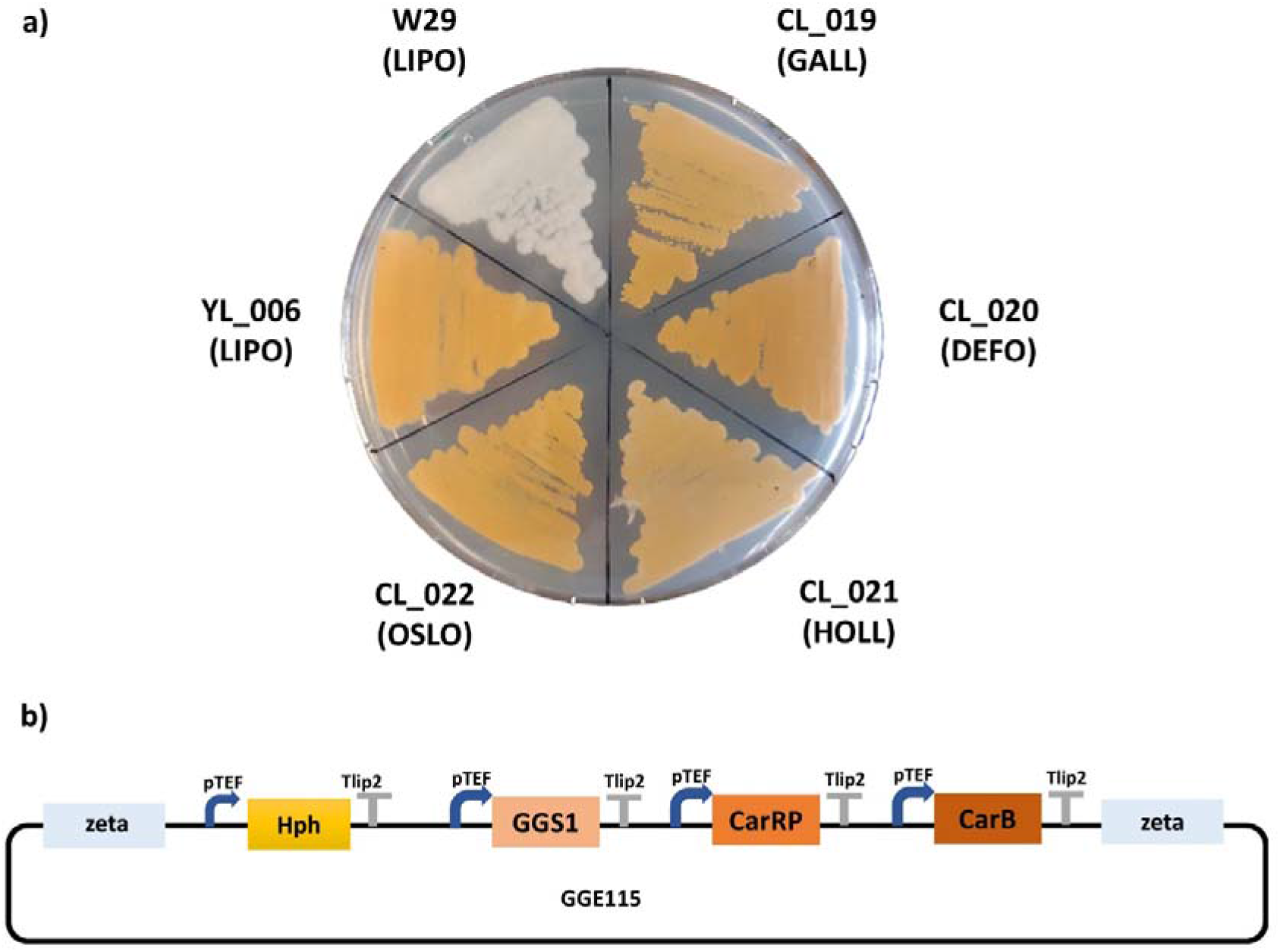
A) Schematic draw of the vector GGE115 used for transformation. B) Growth on YNB plates of strains transformed with the GG115 plasmid for carotenoid pathway expression. LIPO (*Y. lipolytica*); OSLO (*Y. oslonensis*); DEFO (*Y. deformans*); GALL (*Y. galli*); YAKU (*Y. yakushimensis*); HOLL (*Y. hollandica*); W29 (*Y. lipolytica* wild-type strain not transformed). All the strains are listed in supplementary Table 1.

## Conclusion

In this study, we demonstrate that the synthetic biology tools developed for the yeast *Y. lipolytica* can be effectively repurposed for genetic engineering of other members of the clades, in particular for *Y. oslonensis, Y. deformans, Y. galli, Y. yakushimensis* and *Y. hollandica* for which we were able to express a multi-gene heterologous pathway. *Y. phangngensis* and *C. hispaniensis* exhibit a distinct GC content compared to other members of the clades and significant reduction in size for some protein families (Groenewald, et al., 2014), indicating a greater sequence divergence. Furthermore, both strains are highly resistant to typical antibiotics, which may suggest lower compatibility, if any, with *Y. lipolytica*’s dedicated tools. On the other hand, *Y. alimentaria* is much closer *to Y. lipolytica* in term of GC content, and its protein expression failure could help determine the homology threshold for compatible expression, especially for promoters, although further investigation is required.

While it is widely recognized that the choice of host is crucial in metabolic engineering applications, there’s still limited understanding of the strain diversity among non-type strains, particularly for non-conventional microorganisms. Therefore, exploring this aspect of diversity is a critical route to achieving success in microbial cell factories. By evaluating other relevant traits, we anticipate that the results presented in this study will facilitate new biotechnological applications involving previously overlooked and under-characterized strains within the Yarrowia clade.

## Material and methods

### Strains and media

All the wild-type yeast strains used in this study are listed in Table 1. All constructed strains are listed in supplementary Table 1.

Chemically competent *Escherichia coli* DH5α cells were used for cloning and plasmid propagation. *E. coli* cells were grown at 37°C with constant shaking on 5 ml of LB medium with ampicillin (100 μg ml^-1^) or kanamycin (50 μg ml^-1^) for plasmid selection. Yeasts were grown on rich medium YPD (10 g l^-1^ of yeast extract, 10 g l^-1^ of Peptone, 10 g l^-1^ of glucose) complemented with hygromycin (200 μg ml) or nourseothricin (500 μg ml^-1^) for the selection of transformants. Solid media were prepared by adding 1.5% agar. For growth on 96 well plates, yeasts were grown on YNB minimal medium containing 10 g l^-1^ glucose, 10 g l^-1^, 1.7 g l^-1^ yeast nitrogen base without amino acids, 5.0 g l^-1^ NH4Cl^-1^ and 50 Mm phosphate buffer (pH 6.8).

### Plasmid construction

Plasmids expressing the Redstar2 fluorescent protein were assembled using GoldenGate method as described in (Larroude, et al., 2022) using hygromycin or nourseothricin marker, pTEF promoter, tLip1-3 terminator, ZETA upstream and downstream integration sequences, Redstar2 gene and pSB1A3 backbone vector. All building parts are described in (Larroude, et al., 2019).

Plasmid GGE115 allowing expression of the carotenoid pathway was described in (Larroude, et al., 2022). It contains the ZETA sequences for random integration, the hygromycin marker, and the three genes involved, GGS1 (geranylgeranyl diphosphate synthase) from *Y. lipolytica*, carPR (phytoene synthase/lycopene cyclase) from *Mucor circinelloides*, and carB (phytoene dehydrogenase) from *M. circinelloides* (Larroude, et al., 2018).

### Promoter sequences alignment

The *Y. lipolytica* pTEF promoter was used to retrieve promoter ortholog from Whole Genome Sequences of the other species (*Yarrowia_osloensis* ULGU01000000; *Yarrowia_deformans* ULGY01000000; *Yarrowia_galli* ULGS01000000; *Yarrowia_yakushimensis* ULGW01000000; *Yarrowia_alimentaria* ULGN01000000; *Yarrowia_hollandica* ULGV000000). Multiple sequences alignment was performed using T-Coffee web server (Di Tommaso, et al., 2011)

### Transformation

EZ-Yeast transformation kit (MP Biomedicals) was used for all yeast transformations following the manufacturer’s instruction. The kit does not require competent cells and allows using colony growing on plates followed by a 30 min transformation step. Plasmids were first digested with restriction enzyme NotI to release the expression cassette and 2 μg of digested DNA was used for transformation. For the release of the expression cassette of plasmid GGE115 the restriction enzyme SfiI was used.

### Growth curves and fluorescence measurements

Yeasts were precultured for 24 h in YNB medium and diluted to an OD_600nm_ of 0.1 in fresh medium and 200 μl was transferred into 96-well microplates. The growth analysis was performed using a microtiter plate reader (Synergy Mx; BioTek). The settings were 28°C and constant shaking with a reading every 30 min for 72 h for OD_600nm_ as well as red fluorescence (excitation 558 nm /emission 586 nm). Fluorescence was expressed as a ratio of relative fluorescence divided by OD. Cultures were performed at least in duplicate.

