## Supplementary file 2 for "Broaden the application of *Yarrowia Lipolytica* synthetic biology tools to explore the potential of Yarrowia clade biodiversity"

### T-Coffee alignment result

#### MSA

*The multiple sequence alignment result as produced by T-coffee.*

```
T-COFFEE, Version_11.00 (Version_11.00)
Cedric Notredame
SCORE=822
*
 BAD AVG GOOD
*
pTEF Yarrowia l     :  72
Yarrowia osloen     :  73
Yarrowia deform     :  74
Yarrowia galli      :  73
Yarrowia yakush     :  71
Yarrowia alimen     :  68
Yarrowia_hollan     :  70
cons                :  82

pTEF_Yarrowia_l     GGGT---------TGGCGGCG-TATTTGTGTCC-CAAAAAACAGCCCCAATTGC-C-CCAATT
Yarrowia_osloen     GGGT-C------GTGGAGGCGATATTTTCGCCCGGCGGAAACA--CACAATTGC-T-CCAATT
Yarrowia_deform     GGT----------TAGAGTCG-CTTAATCGACAATCAAAAACCAT-ACGATTGA-T-CCAATT
Yarrowia_galli      GGGG-T-------TAGAGACT-CAATATTGACAATC-AAAGTGGCCAGATTCGT-T-GCAATT
Yarrowia_yakush     GAGAGT----G-TTTGAGAGA-GAATATTGAGAATGTAAACTCGA-GCAATTGG-T-CATATT
Yarrowia_alimen     GGCC-AAAAGTCGTGCAGCCGGCAATTCCGCCA----ATTGACG--GCAATTGCGT-GCAATT
Yarrowia_hollan     AGAC-G----G-CATGTGGCC-GAAAGCA--GGGCGAAAAACAGT-GAATTCCA-TCCTAATT

cons                                 *                                *         ***


pTEF_Yarrowia_l     G-ACCCCAAATTGA----CCCAGTAGCGGGCCCAACC-CCG-GCGAGAGCCCCCTTCACCCCA
Yarrowia_osloen     G-GCGCCCAATTGA----CCCTGTAAACTCCCCAGAGCCCC-ACACCC-GCTCCCTCACCCCA
Yarrowia_deform     G-ACGTGCAATTGA----CCGGGTGGCATTTTCAGAGTCTC-ACCTTA-CCTCCCAC-CCCCA
Yarrowia_galli      G-ACCTGCAATTGA----CCCTGTAGCTTCTCCAGCTCGCC-ACCACC-CCTACATCACCCCA
Yarrowia_yakush     GCGCCCACAATTGA----CCCTACACGTCTCTCGCAGCCCC-AGAACC-ACCCAATGACCCCA
Yarrowia_alimen     G-GACGGCATTTTGCACGCCCAGCCGCCTCCCCACGTTGCCTACAACC--CCACGAGTCCCGC
Yarrowia_hollan     G-AGCACCAATTGA----CTCTGTGAGATCCTGAACCACTC-ATATCA--CACCCACACCCCA

cons                *       * **      *                               *       ***  


pTEF_Yarrowia_l     CATATCAAAC--CTCCCCCGGTTCCCACACTTGCCGTTAAGGGCGTAGGGTACTGCAGTCTGG
Yarrowia_osloen     CACAACTCGT--CTCCCCCGGTTCTCACACTTGCCGTTAAGGGCGTAGGGTACTGCAGTCTGG
Yarrowia_deform     CAAAACCCCG--TTTCCCCCACCGTCACACTTGCCGTTAAGGGCGTAGGCAGCTGCAGTCTGG
Yarrowia_galli      CACAACACTG--TTCCCCCAGTTCTCACACTTGCCGTTAAGGGCGTAGGCGTCTGCAGTCTGG
Yarrowia_yakush     CACAACTCTC--TTGCCCCAGTTGTCACACTTGCCGTTAAGGGCGTAGAGTACTGCAGTCTGG
Yarrowia_alimen     CACCTCTGGGGTCGCTATTGGGGCTTGAGTTTCCCGTT-AGGGCGTAGGGTAATGCAGTCGGT
Yarrowia_hollan     CACTCAGCAC--TCTCCCCGGCACTTCAGTTTCCCGTTAAGGGCGTAGGGTACTGCAGTCTGG

cons                **                            ** ***** *********     ******* * 


pTEF_Yarrowia_l     AATCTACGCTTGTTCAGACTTTGTACTAGTTTCTTTGTCTGGCCATCCGGGTAACCCATGCCG
Yarrowia_osloen     AATCTACGCTTGTTCAGACTTCGGGCCGGTTTCTTTGTCCAGCCATCCGGGTAACCCATGCCG
Yarrowia_deform     AATCTACGCTTGTTCAGAATTCGGACCTTTTTCTTTGTCCGGCCATCCGGGTAACCCATGCCG
Yarrowia_galli      AATCTACGCTTGTTCAGAATTCAGCCCTGTTTCTTTGTCCAGCCATCCGGGTAACCCATGCCG
Yarrowia_yakush     AATCTACGCATGTTCAGACTTTGGCACTGTTTCTTTGTCTGACGATCCGGGTAACCCATGCTG
Yarrowia_alimen     AAGCTACATCCATACAGAGTTTAGTCTCCTTTCTTTGTCCGCTAGGTCGGGTAACCCATGCTT
Yarrowia_hollan     AACCTACGATCGTTCAGAGTTTAGACCACTTTCTTTGTCTGACACTCCGGGTAACCCATGCTG

cons                ** ****     * **** **        **********        **************  


pTEF_Yarrowia_l     GACGCAAAATAGACTACTGAAAATTTTTTTGCTT-TGTGGTTGGGACTTTAGCCAAGGGTATA
Yarrowia_osloen     GACGCAAAATAGACTACTGAAAATTTTTTTGCTT-TGTGGTTGGGACTTTAGCCAAGGGTATA
Yarrowia_deform     ACCGCAAAATAGACTACTGAAAATTTTTATGCTC-TGTGGTTGGGAGGTAGCCCCAGGGTATA
Yarrowia_galli      GACGCAAAATAGACTACTGAAAATTTTTATGCTC-TGTGGTTGGGACTTTAGCCAAGGGTATA
Yarrowia_yakush     GGCGGCAAATAGCCTTTTGAAAATTTTTTTGCCT-GGAGGTTGGGACTTTGGCCCAGGGTATA
Yarrowia_alimen     GTCGCT-TTTAGAGTCATGAAAATTTTTTTTTTGCAACACCTGCGTCTTCGCTCCCAGGTATA
Yarrowia_hollan     GGTGCAAAATAGACTACTGAAAATTTTTTTGCTT-TGTGGTTGGGACTTGAGCCAAGGGTATA

cons                   *     ***  *  *********** *           ** *   *    *   ******


pTEF_Yarrowia_l     AAAGACCACCGTCCCCGAATTACCTTTCCTCTTCTTTTCTCTCTCTCCTTGT-CAACTCACAC
Yarrowia_osloen     AAAGACCACCGTCCCCGAATTACCTTTCCTCT---TTTCTCTCTCTCCTTGT-CAACTCACAC
Yarrowia_deform     AAAGACCACCGTCCCCGAATTACCTTTCCTCTTCTTTTCTCTCTCTCCTTGT-CAACTCACAC
Yarrowia_galli      AAAGACCACCGTCCCCGAATTACCTTTCCTCTTCTTTTCTCTCTCTCCTTGT-CAACTCACAC
Yarrowia_yakush     AAAGACCACCGTCCCCGAATTACCTTTCCTCTTCTTTTCTCTCTCTCCTTGTATTAGTTTCAT
Yarrowia_alimen     AAAGAGACGAGTTTCCGAATTACCTTTCTTCTTTTTTTCT---TCTCCTTGT-CAACTCACAC
Yarrowia_hollan     AAAGACCACCGTCCCCGAATTACCTTTCCTCTTCTTTTCTCTCTCTCCTTGT-CAACTCACAC

cons                *****     **  ************** ***   *****   *********   * *  ** 


pTEF_Yarrowia_l     CCGAA
Yarrowia_osloen     CTGAA
Yarrowia_deform     CCGAA
Yarrowia_galli      CTGAA
Yarrowia_yakush     -TCAA
Yarrowia_alimen     CTGAA
Yarrowia_hollan     CCGAA

cons                   **
```
