## Supplementary table 1 for "Broaden the application of *Yarrowia Lipolytica* synthetic biology tools to explore the potential of Yarrowia clade biodiversity"

Supplementary table 1. List of strains generated in this study

| Strain number | specie | Vector used |
| --- | --- | --- |
| YL_001 | Yarrowia lipolytica W29 | HGH pTEF-RS2 |
| YL_005 | Yarrowia lipolytica W29 | NAT pTEF-RS2 |
| CL_004 | Candida oslonensis CBS 10146 | -HGH-pTEF RS2 |
| CL_006 | Yarrowia deformans CBS 2071 | -HGH-pTEF RS2 |
| CL_008 | Candida galli CBS 9722 | -HGH-pTEF RS2 |
| CL_010 | Yarrowia yakushimensis CBS 10253 | -HGH-pTEF RS2 |
| CL_017 | Candida hollandica CBS 4855 | -HGH-pTEF RS2 |
| CL_001 | Candida oslonensis CBS 10146 | -NAT-pTEF RS2 |
| CL_011 | Yarrowia deformans CBS 2071 | NAT-pTEF RS2 |
| CL_013 | Candida galli CBS 9722 | NAT-pTEF RS2 |
| CL_015 | Yarrowia yakushimensis CBS 10253 | NAT-pTEF RS2 |
| CL_003 | Candida hollandica CBS 4855 | -NAT-pTEF RS2 |
| Y8750 CL_019 | Candida galli CBS 9722 | GGE115 |
| Y8753 CL_020 | Yarrowia deformans CBS 2071 | GGE115 |
| Y8755 CL_021 | Candida hollandica CBS 4855 | GGE115 |
| Y8756 CL_022 | Candida oslonensis CBS 10146 | GGE115 |
| Y8760 YL_006 | Yarrowia lipolytica W29 | GGE115 |
